## Supplementary material for "Does protein A mirror image exist in solution? Outline of an experimental design aimed to detect it": suplemental file

Supplementary files for: **Does protein A mirror image exist in solution? Outline of an experimental design aimed to detect it**

Osvaldo A. Martin[1], Yury Vorobjev[2], Harold A. Scheraga[3], and Jorge A. Vila[1,3]

1 IMASL-CONICET, Universidad Nacional de San Luis, Ejército de Los Andes 950, 5700. San Luis, Argentina
2 Institute of Chemical Biology and Fundamental Medicine of the Siberian Branch of the Russian Academy of Science, Lavrentiev Avenue 8, Novosibirsk 630090, Russia
3 Baker Laboratory of Chemistry and Chemical Biology, Cornell University, Ithaca, NY 14853-1301

**Table 1**. pKa constants for the initial native-like and mirror-image structures of Q10H protein.

| Residue | Native-like | Mirror-image |
|---|---|---|
| NEND | 8.2 | 7.9 |
| ASP3 | 3.6 | 2.9 |
| LYS5 | 10.8 | 10.9 |
| LYS8 | 10.7 | 12.4 |
| GLU9 | 2.6 | 3.6 |
| **HIS10** | **5.8** | **7.3** |
| TYR15 | 9.5 | 9.5 |
| GLU16 | 4.2 | 3.7 |
| HIS19 | 7.1 | 7.0 |
| GLU25 | 3.6 | 3.4 |
| GLU26 | 3.8 | 3.8 |
| LYS36 | 11.1 | 10.9 |
| ASP37 | 3.5 | 3.7 |
| ASP38 | 3.1 | 3.3 |
| GLU48 | 3.4 | 3.2 |
| LYS50 | 11.8 | 10.9 |
| LYS51 | 11.6 | 11.3 |
| ASP54 | 2.6 | 3.1 |
| LYS59 | 9.9 | 9.9 |
| CEND | 3.9 | 3.8 |

**Table 2**. Low energies ionization states of native-like and mirror-image structures of Q10H protein.

| protein | Ionization state[a] | $E_{pot}$[b] | RMSD[d] |
|---|---|---|---|
| native | 22 | 88.6 | 24.3 |
| | 20 | 100.1 | 25.6 |
| | 21 | 104.7 | 29.3 |
| Mirror-image | 22 | 183.8 | 24.5 |
| | 20 | 184.0 | 24.5 |
| | 02 | 188.7 | 24.1 |
| | 00 | 189.0 | 23.1 |

[a] Ionization state of histidine residues H10 and H19, where 0,1,2 are integers designating the H+, the HID and the HIE forms, respectively. Thus, for example, the ionization state 20 means that H10 and H19 are in the HIE and H+ forms, respectively.
[b] Average potential energy along the pH-constant 25ns MD trajectory, kcal/mole ;
[c] RMSD energy-fluctuation, at 300K, along the pH-constant 25ns MD trajectory, kcal/mole.

**Table 3**. Average pKa constant and fractions of the His10 forms on the native-like and mirror-image conformations along the 25ns constant-pH MD simulation at pH 7.0.

| Protein | $\langle E_{pot} \rangle$[d]<br>RMSD[e] | His10[f]<br>$\langle pK \rangle$[g] | H+[h] | HID[i] | HIE[j] | His19[f]<br>$\langle pK \rangle$[g] | H+[h] | HID[i] | HIE[j] |
|---|---|---|---|---|---|---|---|---|---|
| Mirror-initial[a] | n/a | 7.6 | 0.76 | 0.08 | 0.16 | 7.1 | 0.58 | 0.14 | 0.37 |
| Mirror-image[b] | 183.9<br>(24.2) | **7.30**<br>(0.21) | 0.65<br>(0.08) | 0.07<br>(0.02) | 0.28<br>(0.07) | 7.00<br>(0.21) | 0.49<br>(0.03) | 0.10<br>(0.01) | 0.41<br>(0.03) |
| Native-like[c] | 88.6<br>(24.3) | **6.20**<br>(0.27) | 0.15<br>(0.06) | 0.09<br>(0.02) | 0.76<br>(0.08) | 7.03<br>(0.19) | 0.51<br>(0.06) | 0.08<br>(0.01) | 0.41<br>(0.06) |

[a] Initial mirror-image structure;

[b] MD optimized  mirror-image structure;

[c] MD optimized native-like structure;

[d] Average potential energy, kcal/mole;;

[e] RMSD due to energy fluctuations (in parenthesis), kcal/mole;;

[f] Histidine residue number;

[g] Average pK and, in parenthesis, its RMSD due to fluctuations;

[h] Average fraction of ionization states and, in parenthesis, its RMSD due to fluctuations;

[i] Average fraction of HID tautomer and, in parenthesis, its RMSD due to fluctuations;

[j] Average fraction of HIE tautomer and, in parenthesis, its RMSD due to fluctuations;

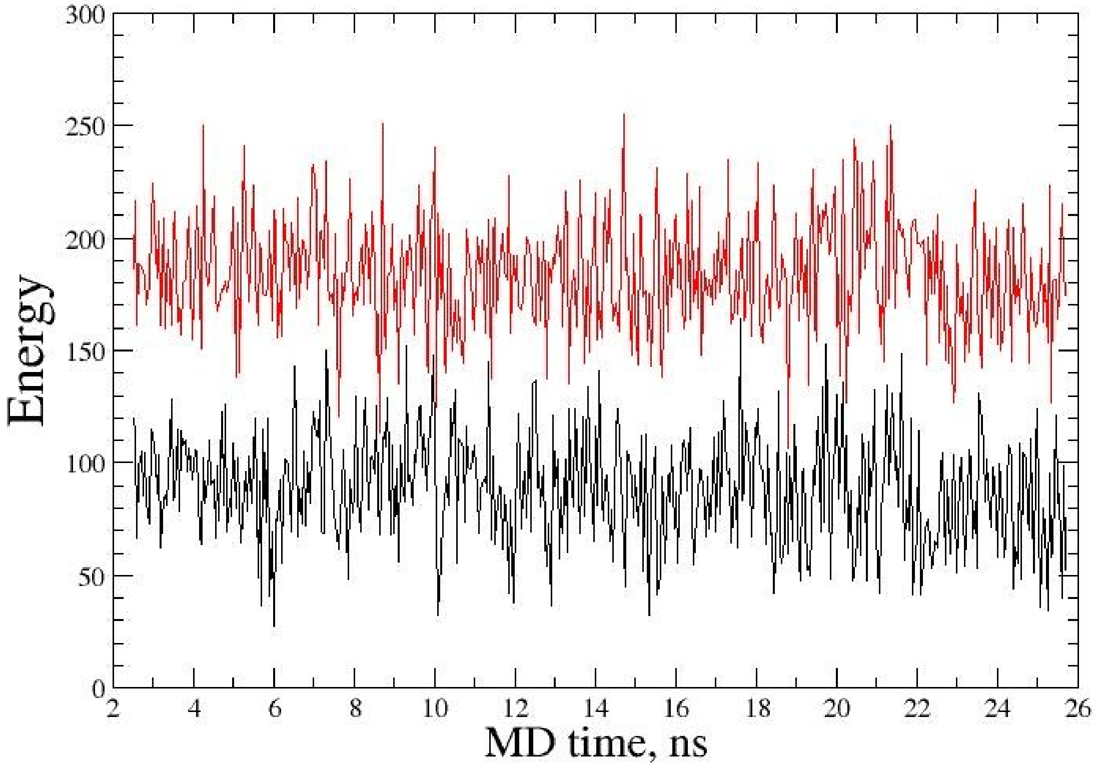


**Figure 5**. Potential energies for *native-like* and *mirror-image* structures of mutant Q10H of protein A (PDB ID 1BDD) along MD trajectories of 25ns; black – native-like; red – mirror-image.

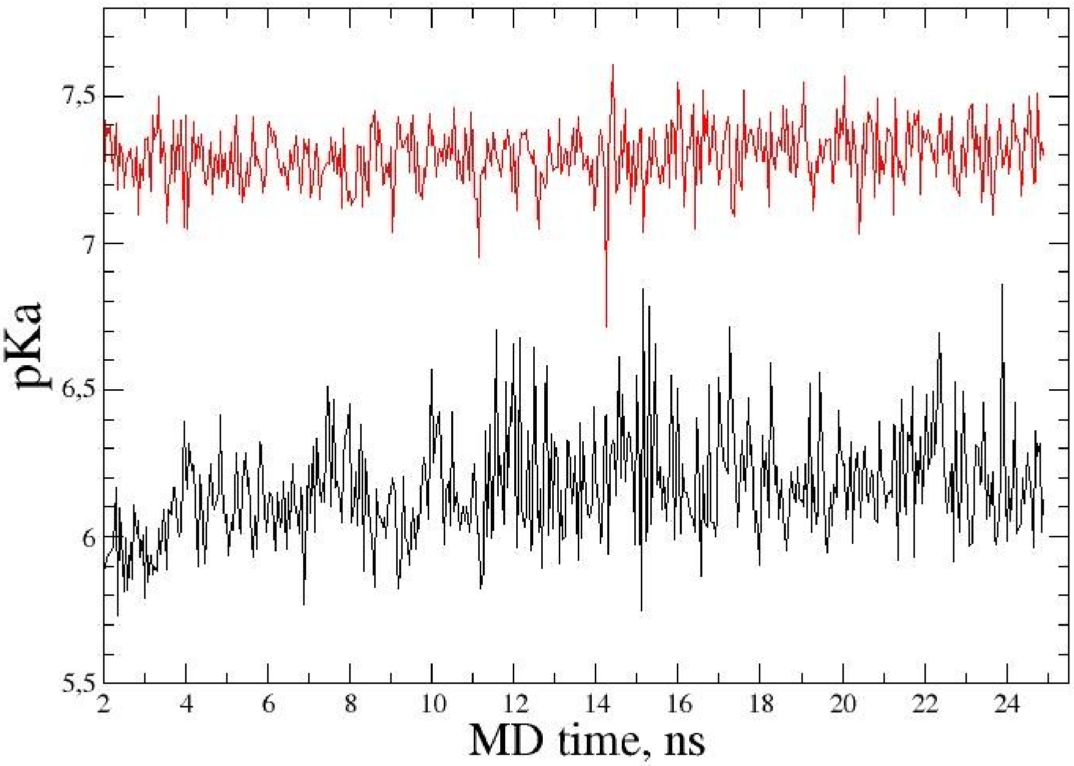


**Figure 6**. Ionization constant pKa of *native-like* and *mirror-image* structures of mutant Q10H of protein A (PDB ID 1BDD) along MD trajectories of 25ns; black – native; red – mirror-image.

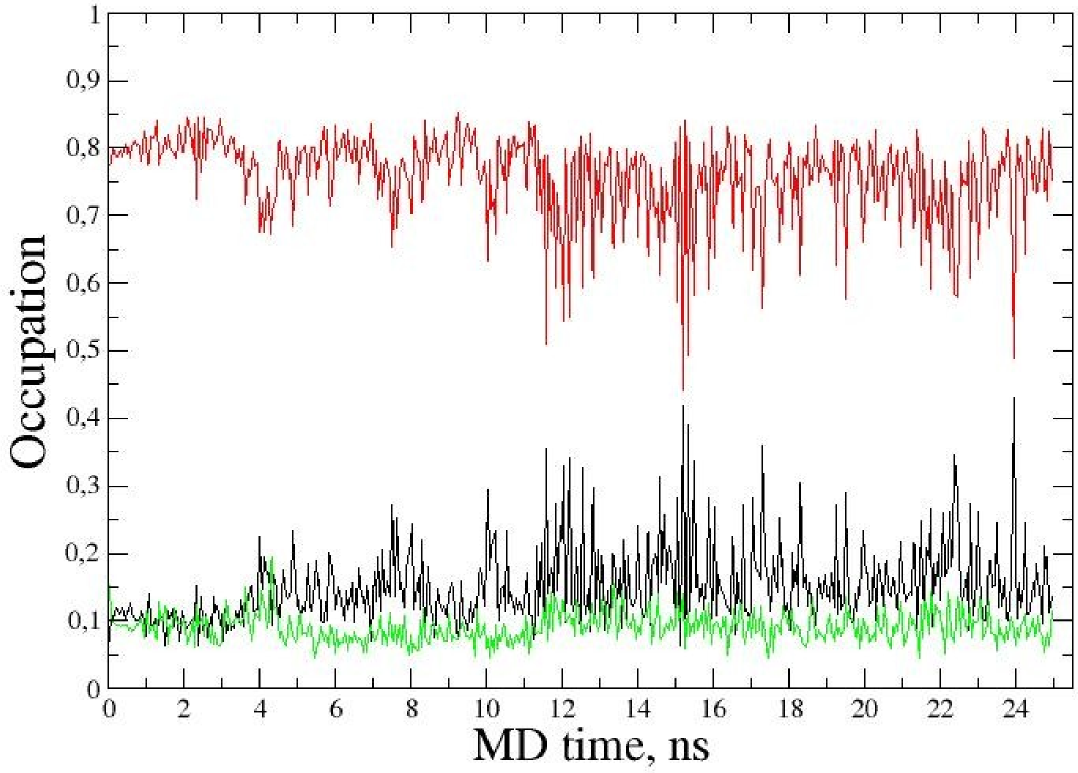


**Figure 7a.** Occupation of ionization states of histidine His10 for *native-like* structure of mutant Q10H; black – ionized state; red – HIE tautomer; green – HID tautomer.

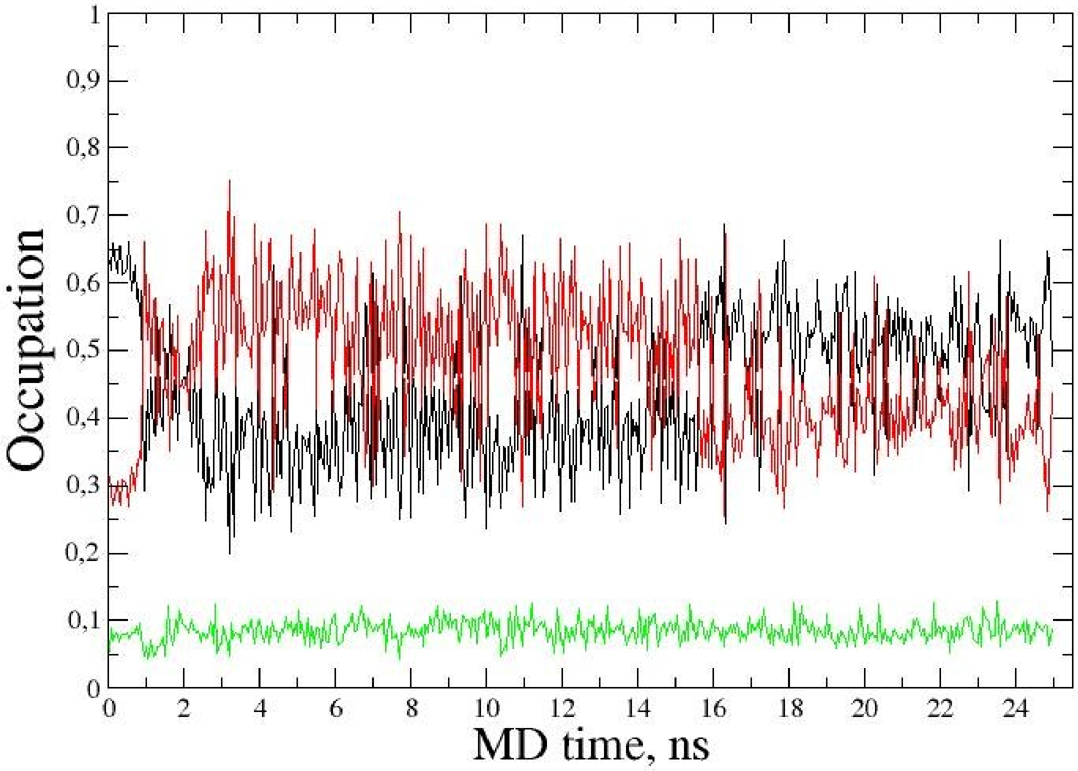


**Figure 7b**. Occupation of ionization states of histidine His10 for *mirror-image* structure of mutant Q10H; black – ionized state; red – HIE tautomer; green – HID tautomer.
